# Switching to external flows: perturbations of developing vasculature within chicken chorioallantoic membrane

**DOI:** 10.1101/2024.01.11.575208

**Authors:** Prasanna Padmanaban, Danny van Galen, Nasim Salehi-Nik, Mariia Zakharova, Loes Segerink, Jeroen Rouwkema

## Abstract

The impact of fluid flow shear stresses, generated by the movement of blood through vasculature, on the organization and maturation of vessels is widely recognized. Nevertheless, it remains uncertain whether external fluid flows outside of the vasculature in the surrounding tissue can similarly play a role in governing these processes. In this research, we introduce an innovative technique called superfusion-induced vascular steering (SIVS). SIVS involves the controlled imposition of external fluid flow patterns onto the vascularized chick chorioallantoic membrane (CAM), allowing us to observe how this impacts the organization of vascular networks. To investigate the concept of SIVS, we conducted superfusion experiments on the intact chick CAM cultured within engineered eggshell system, using phosphate buffered saline (PBS). To capture and analyze the effects of superfusion, we employed a custom-built microscopy setup, enabling us to image both superfused and non-superfused regions within the developing CAM. This study provides valuable insights into the practical application of fluid superfusion within an *in vivo* context, shedding light on its significance for understanding tissue development and manipulation in an engineering setting.

## Introduction

Physical transport mechanisms, including oxygen transport, nutrients distribution, and gas exchange facilitated by flowing liquid within vascular networks and the surrounding interstitial spaces are widely acknowledged as critical for tissue development and for shaping the overall architecture of vascular networks within tissues.^1,2^ These flow processes result in the generation of varying shear stresses within multi-sized vessels, which are governed by the pulsating heart in living organisms, as investigated using models such as chick^3^, zebrafish^4^, and frog embryos.^5^ Conversely, within developing vessels of engineered tissues in for instance microfluidic chips, these shear stresses are regulated by the pressure driven flows generated by micropumps. The former process involves complex flow patterns which are challenging to control, whereas the latter can be modulated by adjusting the flowrates or applied pressure gradients.

Complex flow patterns arising within intricate vascular geometries and interstitial tissue space include laminar, pulsatile^6^, oscillatory^7^, and turbulent profiles.^8^ It is known that fluid flow shear stresses arising from flows within vessels direct the organization and maturation of the vasculature.^9,10^ Previous studies have demonstrated that the development and organization of vascular networks and associated flows during embryogenesis can be altered through genetics and chemical treatments.^11,12^ Moreover, these vascular networks exhibit adaptability in response to hemodynamics and can be highly influenced by the environment.^9,13^ However, it has not been determined whether external fluid flows, which are generally easier to control in an engineering setting, can also influence vascular networks. Therefore, to exert localized control over fluid flow induced shear stresses in vivo, we aim to explore methods that extend beyond genetic and chemical interventions and to actively direct the development and organization of vasculature using the support of controlled externally induced flows.

In this manuscript, we demonstrated an innovative strategy termed superfusion-induced vascular steering (SIVS), which involves the controlled application of external fluid flow patterns onto the vascularized CAM tissue **(Figure 1)**. We accomplished this by utilizing an engineered eggshell platform to culture the entire chick CAM specimens, which possess multi-sized vasculature networks **(Supplementary Video 3a)**. Additionally, we integrated a perfusable microfluidic channel designed to facilitate the fluid flow of varying viscosities onto the surrounding microenvironment of the CAM tissue **(Figure 2C-D and Supplementary Video 3b)**. Using this methodology, we introduced changes to the CAM tissue by perfusing PBS and closely examining how this impacts the vascular organization in various regions of the CAM tissue **(Figure 4C-D, Supplementary Videos 4a and 4b)**. As a proof-of-concept test, we inserted a porous polydimethylsiloxane (PDMS) membrane into the microfluidic channel to further enhance the localized, controlled influence of fluidic shear stress on the adjacent microenvironment **(Figure 5A-B and Supplementary Figure S3 and Supplementary Video 5)**.

**Figure 1:**
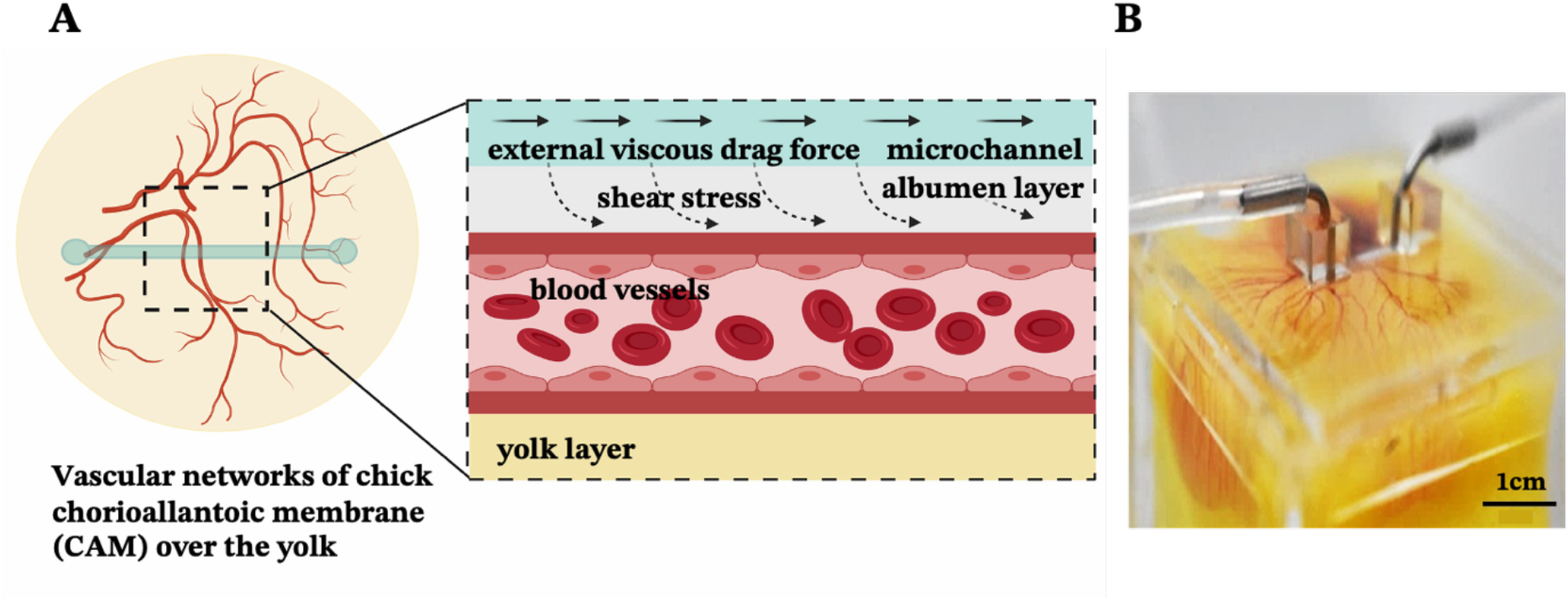
Superfusion induced vascular steering concept. **(A)** represents the illustration of external flow perturbations within developing vascular networks of chick chorioallantoic membrane (CAM) **(B)** shows the experimental sample with microchannels integrated within developing chick CAM cultured in engineered eggshell system.

**Figure 2:**
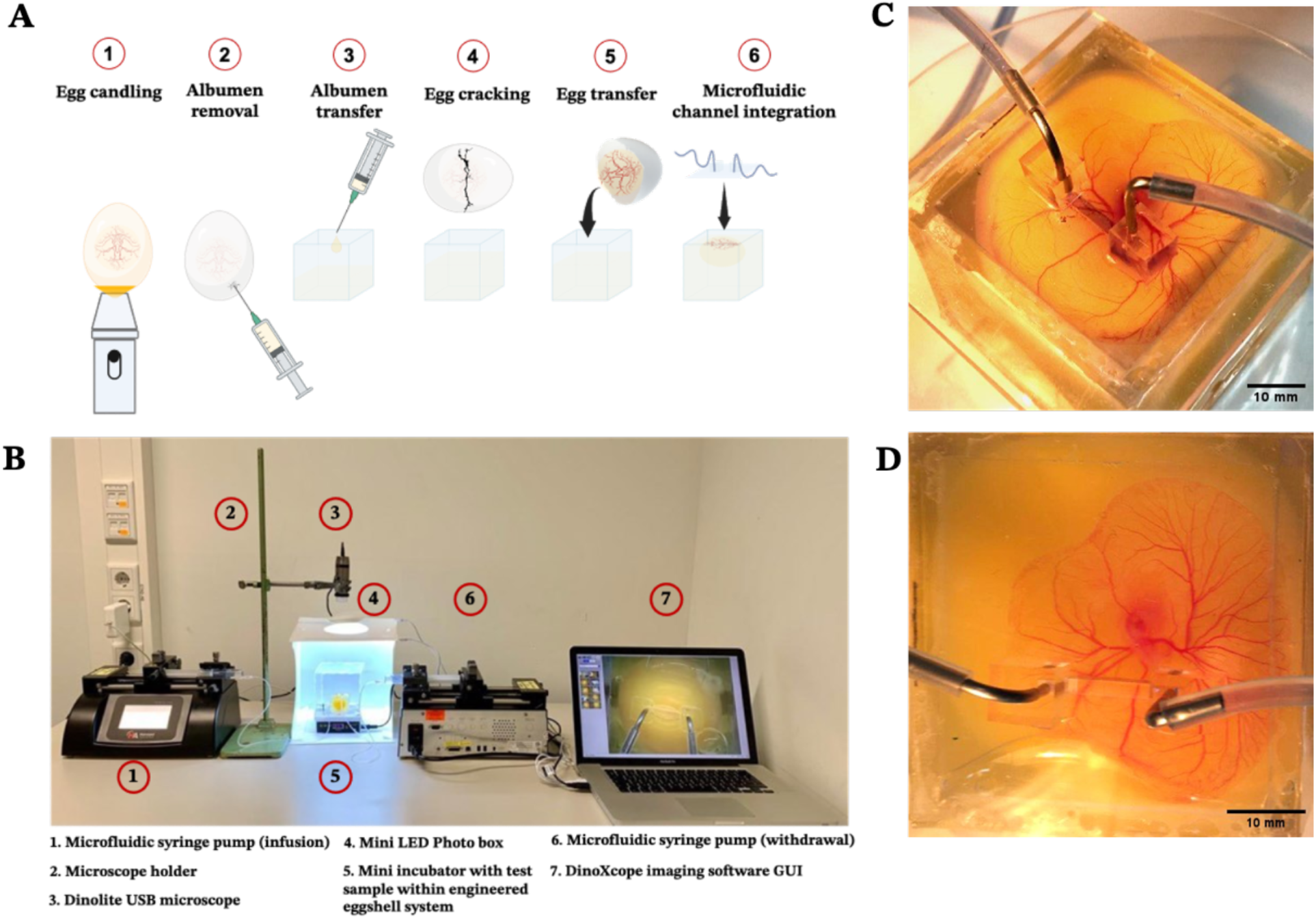
Experimental procedure and microscopy setup for in vivo superfusion. **(A)** depicts an illustration of the experimental protocol for culturing chick CAM using the engineered eggshell system and the incorporation of microfluidic channels **(B)** demonstrates the complete custom microscopy setup and the configuration of microfluidic superfusion system **(C-D)** exhibits the experimental test samples, showcasing the placement of microfluidic superfusion channels at various locations on top of the CAM tissue.

**Figure 3:**
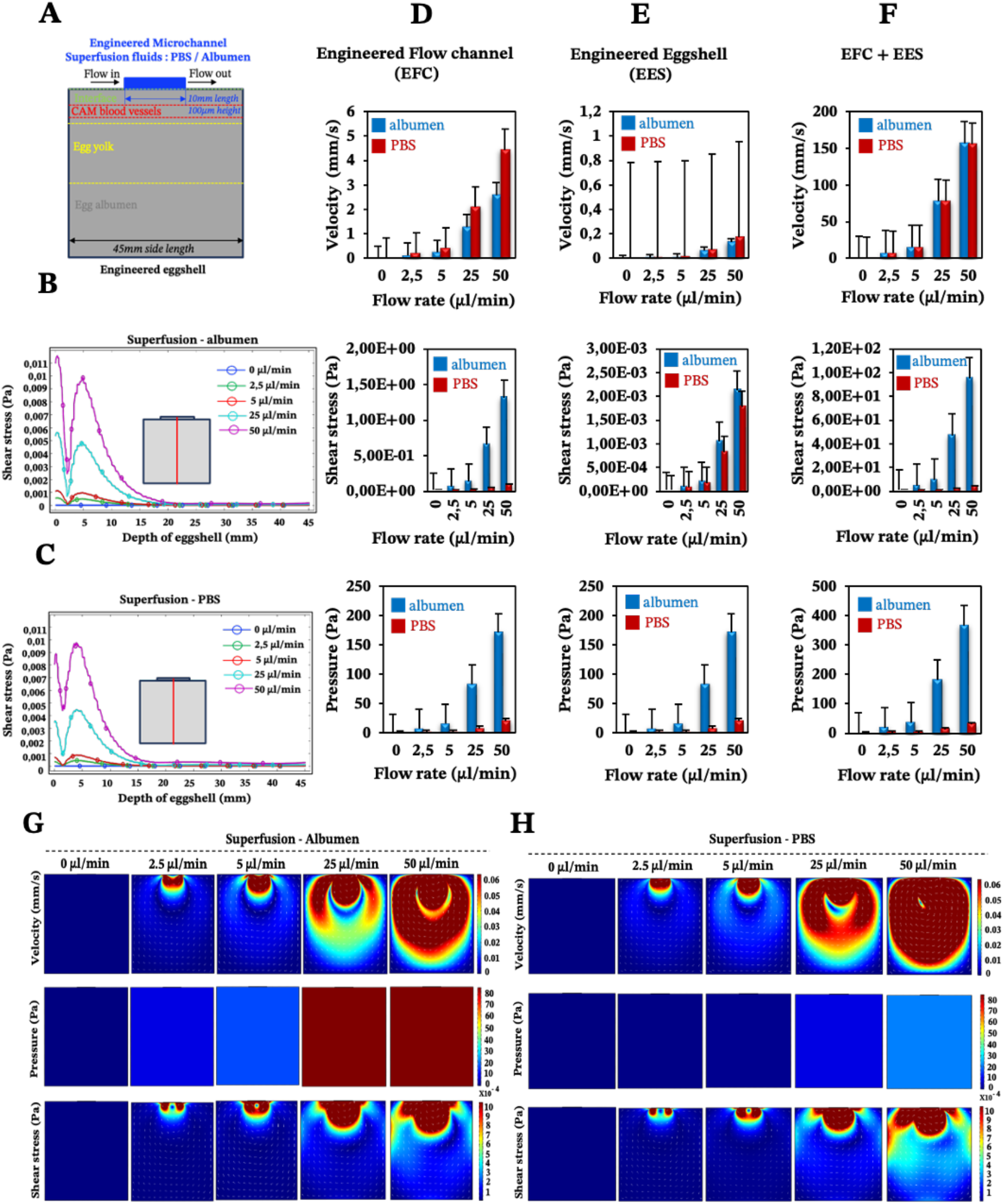
Impact of superfusion on shear stresses and velocities in the engineered eggshell system. This figure portrays 2D simulation data showcasing microfluidic superfusion in an engineered eggshell system. **(A)** presents an illustration of 2D cross sectional view of engineered eggshell, featuring the superfusion channel that served as the geometrical input for the simulation. It also outlines the locations of microchannel, interface, CAM blood vessels, egg yolk and albumen respectively. **(B-C)** indicate the line plot of shear stresses along the depth of engineered eggshell at increasing flow rates. Red line in the cartoon highlights the location of a cut line made in 2D model for analysis. **(D-F)** show the averaged simulated values of velocities, pressure, and shear stresses in two compartments: the engineered flow channel (EFC) and the engineered eggshell (EES), as well as both EFC and EES. This involves superfusing albumen and PBS fluids through integrated microchannels at progressively increasing flow rates. **(G-H)** show the simulation predictions within the engineered eggshell upon albumen and PBS superfusion at increasing flow rates. Heatmaps were plotted using COMSOL Multiphysics software Version 5.6 and graphs were plotted using Microsoft Excel software application.

**Figure 4:**
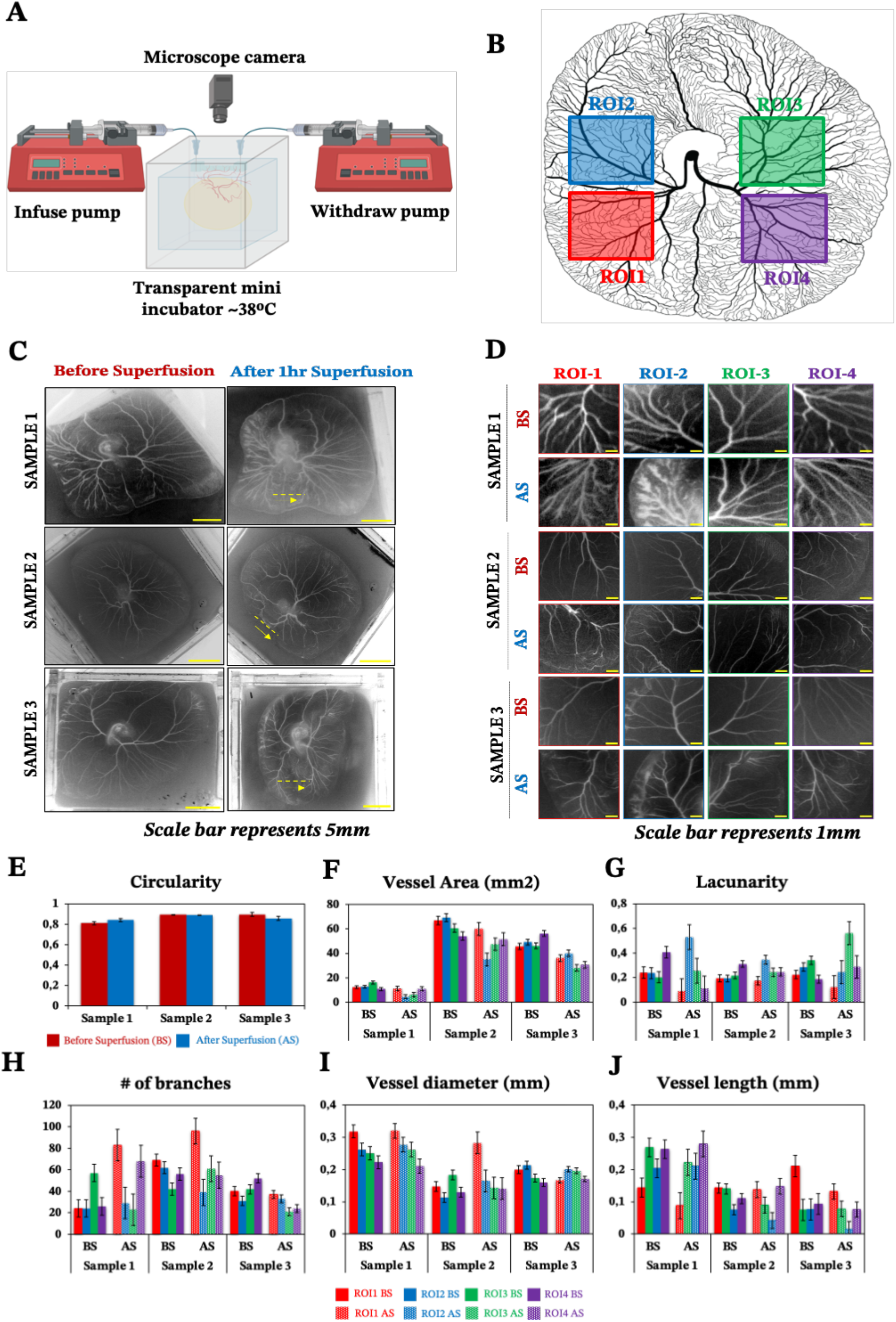
Impact of superfusion on the organization of vascular networks within CAM. This figure portrays experimental observations of 3 different test samples. **(A)** illustrates the schematic representation of the perfusion setup **(B)** shows the selected regions of interest for data analysis **(C-D)** highlight the spatial distribution of vascular networks of CAM tissue cultured in engineered eggshell system before and after the superfusion of PBS at a flow rate of 25 μL/min. Note, the yellow dashed line and arrow line represent the location of microchannel placement and superfused flow direction **(E-J)** represents the quantitative morphometric analysis of CAM circularity, vessel area, number of branches, lacunarity, vessel length and vessel diameter respectively BS = before superfusion, AS = after superfusion. Data analyses were performed using Fiji, Angiotool and graphs were plotted using Microsoft Excel software application.

**Figure 5:**
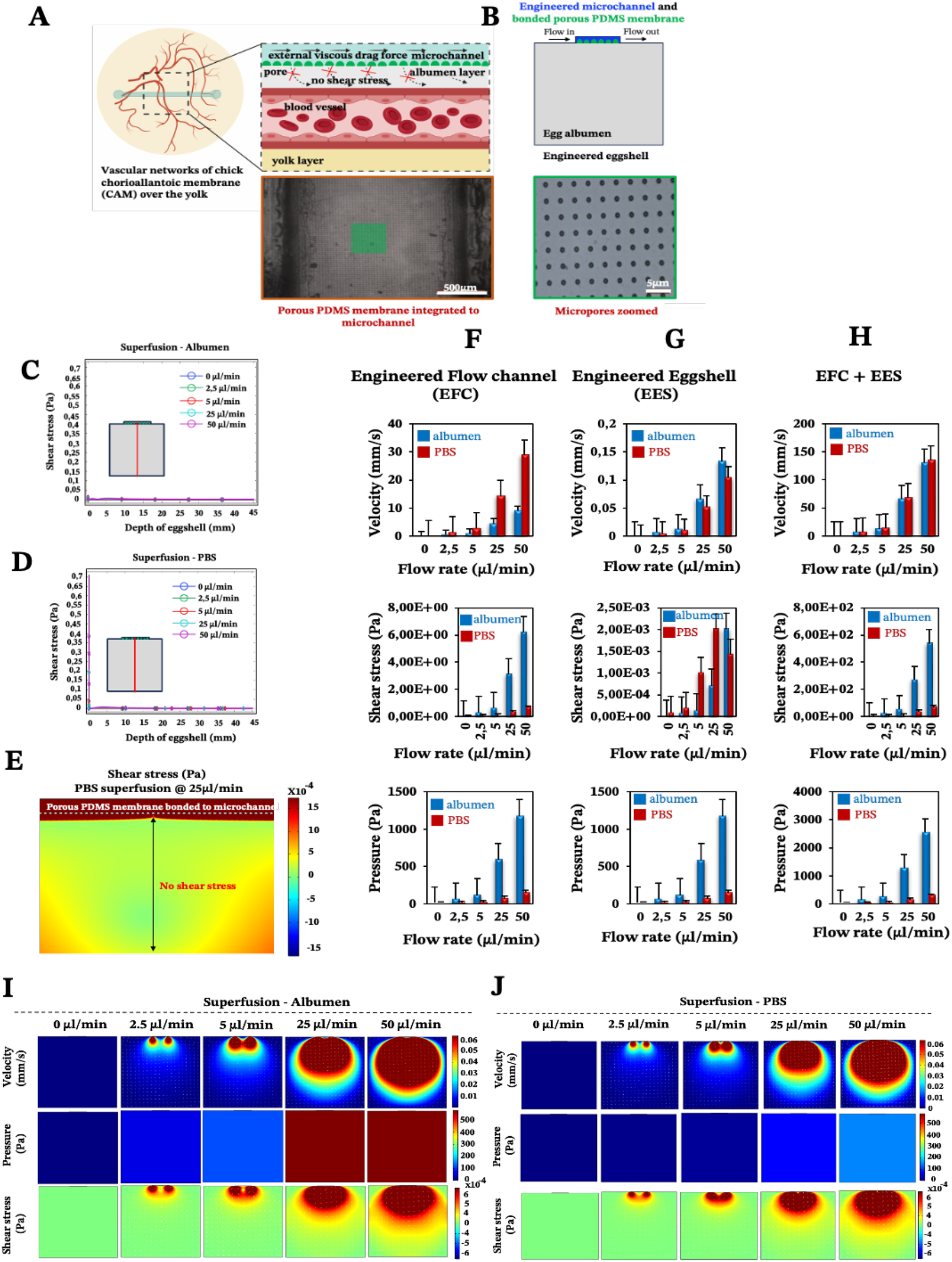
Proof-of-concept study: Impact of porous membrane on superfusion. This figure portrays 2D simulation data of microfluidic superfusion of an engineered eggshell system with a porous membrane between the perfusion channel and the eggshell. **(A)** illustrates the schematic representation of decreased flow perturbations in the presence of a porous membrane **(B)** presents an illustration of 2D cross sectional view of engineered eggshell, showcasing the integrated flow channel with bonded porous membrane served as geometrical input for simulation. Note the bottom panel of A and B show the bonded porous PDMS membrane to the flow channel **(C-D)** indicate the line plot of shear stresses along the depth of engineered eggshell at increasing flow rates. Red line in the cartoon highlights the location of a cut line made in 2D model for analysis. **(E)** highlight 2D model and predicted shear stress distribution within engineered flow channel, interface and engineered eggshell at the location of CAM blood vessels in presence of bonded porous membrane. **(F-H)** emphasize the averaged simulated values within the engineered eggshell upon albumen and PBS superfusion. **(I-J)** display predicted velocities, pressure and shear stresses within the engineered eggshell using albumen and PBS fluids superfused through the integrated microchannels bonded with porous membrane at increasing flow rates. Heatmaps were plotted using COMSOL Multiphysics software Version 5.6 and graphs were plotted using advanced Microsoft Excel software application.

## Results

### Engineered eggshell system with integrated microchannel for in vivo superfusion

The design of the engineered eggshell platform is based on a previously published system that has been adopted and re-engineered for the purpose of this study.^14^ The platform used comprises of two components: (1) a 2mm thick polymethylmethacrylate (PMMA) frame that offers structural support, and (2) thin PDMS membranes with a thickness between 500-700μm, which are employed to create accessible windows as shown in **Figure 1B, Supplementary Figure S1 and S2**. To expedite and streamline the assembly of the eggshell, 3D drawing was generated using the “Lego-cuts” method through www.makercase.com. This approach allows for a quick and efficient construction of the eggshell. For the frame, 2mm thick PMMA sheets were laser-cut and assembled, and quick-fix super glue (Loctite) was used to seal the gaps between the Lego-cut components, ensuring that no liquid leakage would occur. For the membrane containing the flow channel, PMMA rectangular blocks measuring 10mm in length, 1mm in width and 100μm in height were laser-cut and affixed to square-shaped PMMA sheets of 4cm side length, to form a negative mold. The PDMS membrane was subsequently prepared by pouring the PDMS over the mold. After assembly of the system, a 2mm biopsy punch was used to create holes in the microchannel, establishing the inlet and outlet points for connecting the pump tubing as shown in **Figure 1B, 2C and 2D** respectively. The eggs were then introduced and cultured according to the method outlined in the methods section as shown in **Figure 2A**. Using this methodology, chick embryos could be cultured successfully within the engineered eggshell system up to 4 days, after which experiments were terminated for ethical reasons. During this period, CAM development was comparable to in-ovo controls.

### Computational fluid dynamics predictions of superfusion induced fluid flow shear stress within CAM

Computational fluid dynamics (CFD) simulations were employed to estimate the superfusion induced velocities, pressure and shear stresses within engineered eggshell platforms containing an integrated flow channel. A 2D cross sectional view of the integrated microfluidic flow channel within the engineered eggshell, as shown in **Figure 3A**, was designed as the geometric input for the CFD simulation using COMSOL Multiphysics software (Version 5.6). Subsequently, the laminar flow physics module was selected to solve the Navier-Stokes equation, which helped to predict flow velocities, shear stresses, and pressure ranges that were shown as heatmaps in **Figure 3B-H**.

The simulation utilized boundary conditions, where a flow rate-driven model and no-slip conditions were applied to the side walls of the flow channel. At the entrance of the flow channel, inlet flow rates ranging from 0 to 50μl/min were instituted, and a zero-pressure boundary was set at the outlet of the flow channel. The model was subjected to extremely refined physics-controlled mesh conditions consisting of ∼30000 triangular elements. Two different perfusion liquids were simulated; albumen and PBS, represented by densities and viscosities ρ_albumen_= 1035 kg/m^3^, η_albumen_=0.0181 Pa.s; π_PBS_ = 993 kg/m^3^, ρ_PBS_=6.78×10^−4^ Pa.s respectively.^15,16^ As can be seen in **Figure 3**, simulations reveal a direct linear correlation between albumen and PBS superfusion and generated shear stresses with rising flow rates. This unequivocally demonstrates that the viscous drag force generated at the interface (walls of flow channel) directly perturbs the CAM blood vessels positioned beneath, as illustrated in **Figure 3A**.

Owing to the variance in fluid viscosities between albumen and PBS, higher flow rates during superfusion result in PBS flowing more rapidly than albumen. This leads to an increase in flow velocities and a decrease in shear stress levels, as predicted in **Figures 3D-H**, in both the engineered flow channel and engineered eggshell compartments. Moreover, this outcome is reflected in the depth effect within the engineered eggshell, as displayed in **Figure 3B-C**. As illustrated in **Figure 3A and G-H**, the blood vessels within CAM encounter higher shear stresses in comparison to the yolk and surrounding albumen. The higher shear stresses are attributed to the positioning of the flow channel directly over the blood vessels. With elevated flow rates during superfusion, viscous fluids accumulate increased pressure within the engineered eggshell, as shown in the bottom panels of **Figure 3D-F** and middle rows of **Figure 3G and H**.

### Superfusion flow-conditioning alters the organization of developing vascular networks

To study whether superfusion induced fluid flows have an influence on the overall organization of the developing vascular network within the CAM, we carried out superfusion experiments employing a PBS solution. The rationale behind selecting PBS is that it exhibits a viscosity comparable to that of cell culture medium and has minimal interference with the overall growth of the embryo. Following the integration of the microfluidic flow channel on top of the vascularized CAM, we initiated PBS perfusion at a rate of 25μl/min as shown in **Supplementary videos 3b and 4a-b. Figure 4A** depicts the schematic representation of the perfusion setup. As our culture platform allows for comprehensive visualization of the vasculature, multiple regions of interest (ROI; see **Figure 4B**) were chosen for morphometric analysis, using the analysis pipeline as described in the methods section.

Based on our computational analysis, flow velocities of albumen / PBS below 0.2mm/s are predicted at the level of the CAM vasculature during superfusion. This, in turn, results in shear stress level of less than 1 dyn/cm^2^, whereas the physiological tissue flow range during primitive streak formation was reported between 0.1-1.5μm/s at day 1 post incubation.^17^ Superfusion experiments were performed on three individual samples at the fourth day of embryonic development (EDD4). Subsequently, microscopy was conducted both before (BS) and after superfusion (AS) of the entire CAM vascularized tissue to evaluate and compare the superfusion effects. Prior to superfusion, the results depict a noticeable increase in both the vasculature area and the number of branching points over time within four ROIs across three individual samples **Figure 4F and H**. Over the same period, there was a distinct decrease in mean lacunarity, attributed to the increased sprouting of new vessel structures **Figure 4G**. An interesting pattern emerged in relation to vessel diameter and length, suggesting an influence on the structural remodeling process **Figure 4I and J**.

Following superfusion, changes were observed in the organization of the vascular networks; a reduction in the vasculature area corresponding to regression, but surprisingly, an increase in branching points. Vessel diameters appeared to follow a similar pattern as before superfusion, albeit with an increase in size **Figure 4I**. Vessel length exhibited a comparable pattern to the pre-superfusion period **Figure 4J**.

The microfluidic flow channel can not only be used for the superfusion of the CAM with the aim of perturbing the mechanical environment of the CAM, but also as a delivery route for chemical perturbations by perfusing for instance liquids containing specific growth factors. In order to be able to decouple the chemical perturbation from the mechanical perturbation, we performed a proof-of-concept test where a porous PDMS membrane was included between the microfluidic channel and the engineered eggshell as shown in **Figure 5 A** and **B**. Notably, as emphasized in **Figure 5I-J** and **5C-E**, it becomes intriguing to observe that flows and shear stresses within the eggshell diminish, with a particular emphasis on the absence of shear acting on the CAM blood vessels positioned beneath the porous membrane interface.

To comprehensively explore the contributory roles of fluid flow dynamics on vascular development, we conducted a systematic examination of circulating red blood cells in various vessels located in distinct regions of the vascularized CAM tissue of chick embryos. Our initial focus was on the flows directed toward and away from the heart, as these vessels experienced changes in flow patterns and directionality throughout development and remodeling. Additionally, we recorded the circulating red blood cells within multi-sized vessels of arteries, veins and microcapillaries present within CAM tissue **(Supplementary video 1a-b, 2a-c)**. Later, by combining numerical predictions and experimental data, we demonstrate the pivotal role of flow-induced shear stresses in shaping vascular organization throughout development and in response to flow perturbations without and with porous membrane **(Figure 5, Supplementary Figure S3 and Supplementary Video 5)**. This firsthand understanding of shear stress is indispensable for tissue engineers aiming to optimize vascular organization during the perfusion of engineered vascularized tissues.

## Discussion

The development and organization of vascular networks can be influenced in specific spatial ways through genetic, physical, and chemical interventions.^9,10,12,18^ In this context, we introduce the concept of superfusion-induced vascular steering (SIVS) and demonstrate that vascular networks and their associated flow patterns can be adjusted locally through the manipulation of external flows. As far as our knowledge extends, this research represents the first demonstration of applying external flows (superfusion) to vascularized tissue in a live organism. The integration of fluid flow parameters with the configuration of vascular networks advances our comprehensive understanding of structural aspects and mechanical conditions within these developing vascular networks. This holds substantial significance in the regulation of vascular organization.

We engineered an eggshell platform designed to support the development of chick embryos with diverse vascular networks, allowing for the integration of microfluidic flow channels on top of the vascularized CAM tissue. This culture platform offers excellent transparency and accessibility from all sides, owing to its cube design, making it ideal for imaging of the vasculature within the CAM. We recommend our experimental protocol for culturing and transferring the entire egg contents to the engineered eggshell platform (as outlined in **Figure 2A**). However, we acknowledge the associated challenges in the transfer of fertilized egg contents to the engineered eggshell system and the placement of microfluidic flow channels in different regions of the vascularized CAM tissue. First, it is essential to emphasize the significance of pre-loading albumen into the engineered eggshell before cracking the native egg. This precaution minimizes potential harm to the vascular structures within the embryo during the cracking process. Secondly, once the entire egg contents have been transferred to the engineered eggshell system, it is important to remove any excess albumen and any air bubbles that may have formed during the transfer. This step helps prevent the microchannel from slipping out of place after placement. Finally, before initiating perfusion, a gentle thumb pressure should be applied to the microfluidic compartment to ensure secure attachment to the surface of the CAM containing the vasculature.

To delineate the connection between vascular organization, blood flows, and shear stresses, we performed imaging of the overall vascular structure both before and after perfusion in the vascularized CAM tissue. Our microscopy data reveal a notable alteration in the arrangement of vascular networks, including an increase in vessel diameter and branching, which corresponds to a slight decrease in vessel length, area, and lacunarity. Numerical predictions affirm a linear relationship, demonstrating that as external perfusion rates increase, superfusion velocities and the associated shear stresses experienced by the vascular structures in the CAM also increase. This study does not elucidate the biological processes that lead to the vascular remodeling as a response to external fluid flow shear stresses. Even though for capillary structures it is possible that a direct response of endothelial cells to the shear stresses is involved, in larger vessels the endothelial cells are surrounded by supportive smooth muscle cells, which makes a direct response by endothelial cells unlikely. Additionally, the microvessels in the CAM are embedded in a cellularized membrane which will also limit the direct exposure to external fluid flow shear stresses.^19^ Even though it was outside the scope of the present study, a more detailed investigation of the cellular processes involved would help in using the SIVS approach to its full potential.

Moreover, we showed that the addition of a porous membrane to the flow channel reduces the shear stress along the depth of the engineered eggshell. This holds significant interest for tissue engineers, as it opens the possibility of delivering chemical compounds such as growth factors or drugs without the need for exposure to shear stress. The rearrangement of vascular networks can be initiated by external flows and specific geometric constraints. These characteristics lead to a diverse network configuration and flow pattern. We propose that such changes in vascular organization may potentially influence locally generated shear stresses within these networks. We speculate that these changes may contribute to shaping the long-range patterning of vascular networks.

## Conclusion

In conclusion, we demonstrate that the introduction of superfusion during vascular development has the potential to alter the overall organization of vascular networks. We attribute these alterations to the accumulation of shear stresses induced by the viscous drag force present in the walls of microfluidic channels located at the top of developing vasculature. Additionally, we highlight the importance of considering both fluid dynamics and structural changes that embryos undergo in their early stages, as these changes impact the circulating blood flow. Our findings offer a fresh perspective on comprehending and testing the impact of microfluidic flows within an in vivo environment. We anticipate SIVS strategy will inspire further investigations addressing the complexities associated with the incorporation of multidirectional flows within microfluidic chips.

## Acknowledgements

The authors acknowledge the financial support from the ERC consolidator grant (724469) and NWO XS grant (OCENW.XS.021) of Jeroen Rouwkema. Illustrations were made using bio render and Microsoft power point.

## Author contributions

**Conceptualization**: PP, NSM, JR; **Formal Analyses**: PP; **Funding Acquisition:** JR; **Investigation**: PP; **Methodology**: PP, NSM; **Project Administration**: PP, JR; **Software**: PP; **Supervision**: JR; **Validation**: PP, NSM, DG, MZ; **Visualization**: PP, DG; **Original Draft**: PP; **Review and Editing**: PP, NSM, DG, MZ, LS and JR.

## Materials and Methods

### Ethics statement

In accordance with the Dutch animal care guidelines, obtaining IACUC approval for experiments involving chicken embryos is not mandated unless there is an anticipation of hatching. Additionally, IACUC approval is only requisite for experiments involving chick embryos at or beyond EDD14 of development. The embryos utilized in this research were all in the early stages of development, falling between EDD3 and EDD6. The fertilized chicken eggs employed in this study were procured from authorized poultry egg farms in the Netherlands.

### Chick embryo culture

As reported earlier and illustrated in Figure 2A, Dekalb white fertilized chicken eggs were obtained from Het Anker BV in Ochten, The Netherlands, and stored at 12°C.^20^ A Day before egg transfer, the modified incubator was set to 38°C with 65% humidity, which was maintained throughout incubation. In the initial 3-day period, the eggs were rotated every two hours for 15 seconds to prevent embryo adhesion to the eggshell. 4 days after starting the incubation, a small ∼2 mm diameter hole was created in the eggshell with fine tweezers, and a 18G syringe microneedle with plastic syringe was used to withdraw 3mL of albumen to protect the yolk and embryo vasculature during cracking. Carefully, the withdrawn albumen was introduced into the engineered eggshell to minimize the entry of air bubbles. After this, the chicken embryo containing multi-sized vascular networks were transferred to the engineered eggshell in a sterile environment. The PDMS-based lid containing flow channel was subsequently positioned onto the CAM blood vessels. Following careful alignment of flow channel with blood vessels, thumb pressure was gently applied to ensure a secure attachment.

### Computational fluid dynamics simulations

Computational models were employed to determine the distribution of fluid flow shear stresses within the engineered eggshell and the inner walls of microfluidic channel. The shear stress acting upon the albumen layer during perfusion was simulated using COMSOL Multiphysics software (Version 5.6). The Navier-stokes equation was solved under laminar flow conditions, with a no-slip boundary condition assumed for the inner walls of the microchannel.

#### Perfusion setup

As illustrated in Figure 4A, the perfusion experiments were conducted under aseptic conditions, employing two syringe pumps (Harvard Apparatus PHD Ultra) connected to the engineered eggshell. One pump was equipped with a sterile syringe (HSW Norm-Ject 25mL) filled with PBS (Gibco). A sterile 1mm tube (Nordson Medical Tubings) was affixed to the syringe, with the connection point joined to the microchannel inlet on the engineered eggshell. The second pump contained an empty syringe and was similarly connected to the microchannel outlet. Identical flow and withdrawal rates were employed to ensure consistent control of flow.

#### Image Acquisition

Images were captured through continuous video recording using a Dino-Lite USB camera (AM4115 ZT 1.3MP) and an HAYEAR digital microscope (16MP Industrial grade, 150X C-Mount Lens). To produce a time lapse video, the recorded video files were processed using corresponding software: (DinoXcope 2.0 for Mac) and (IC Measure software for Windows). Multiple frames extracted from the processed video were then imported and transformed into 8-bit grayscale images using Fiji software. Vessel morphometrics such as length, diameter and number of branches were quantified using Angiotool plugin.

### PDMS porous membrane fabrication

A thin and transparent porous PDMS membrane was bonded to the microchannels to create a barrier between the flow channels and the chick CAM vasculature. As previously detailed, a 2μm thick, porous PDMS membrane with 5μm pore diameter and 30μm pitch was produced using standard photomask lithography techniques as described in.^21,22^ The PDMS membrane was transferred to the microchannel by first, applying oxygen plasma to bond the surfaces of PDMS as shown in **Figure 5A and B**. Subsequently, the assembly was immersed into the acetone solution, releasing the PDMS membrane from the Si wafer by dissolving the positive photoresist which served as a sacrificial layer. The microchannel with the attached membrane was washed in deionized water and dried under the fume hood.

## Supplementary Videos

**Supplementary video 1a**: Red blood cells flowing through multi-sized venous vessels of vascularized CAM tissue.

**Supplementary video 1b**: Red blood cells flowing through multi-sized micro capillaries of vascularized CAM tissue.

**Supplementary video 2a**: Red blood cells flowing through large artery of vascularized CAM tissue. **Supplementary video 2b**: Red blood cells flowing through artery-veins of vascularized CAM tissue. **Supplementary video 2c**: Red blood cells flowing through artery, microcapillaries and veins of vascularized CAM.

**Supplementary video 3a:** Chick embryo containing hierarchical vascular networks cultured within engineered eggshell system – chick embryo moving within engineered eggshell system.

**Supplementary video 3b:** Chick embryo containing hierarchical vascular networks cultured within engineered eggshell system – integrated microchannel perfusion on CAM vasculature.

**Supplementary video 4a:** External flow application within microchannel integrated on CAM within engineered eggshell system – introducing external fluids via an integrated microchannel.

**Supplementary video 4b:** External flow application within microchannel integrated on CAM within engineered eggshell system – red blood cells moving within capillaries by external fluid perfusion.

**Supplementary video 5:** Proof-of-concept testing of integrated porous membrane to microfluidic channels within engineered eggshell system.

## Supplementary Figures

**S1.**
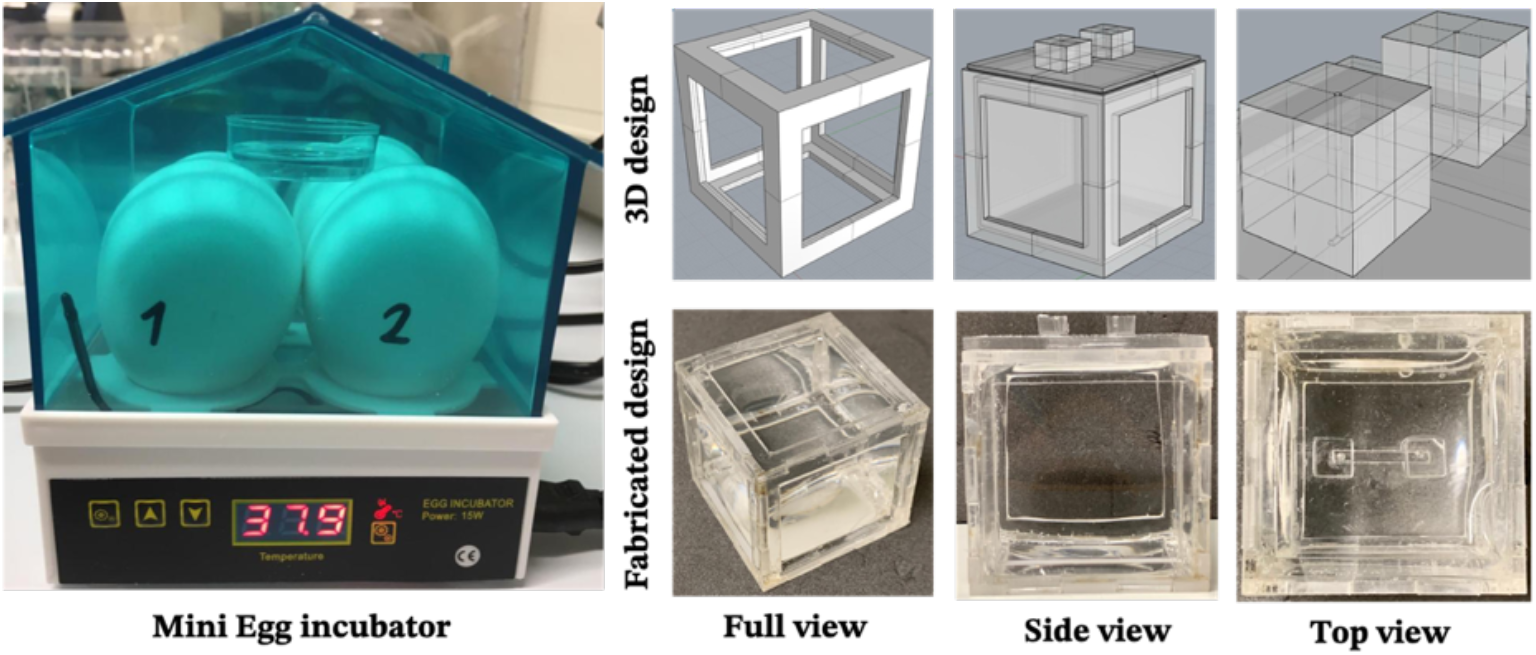
Egg incubator and engineered eggshell design.

**S2.**
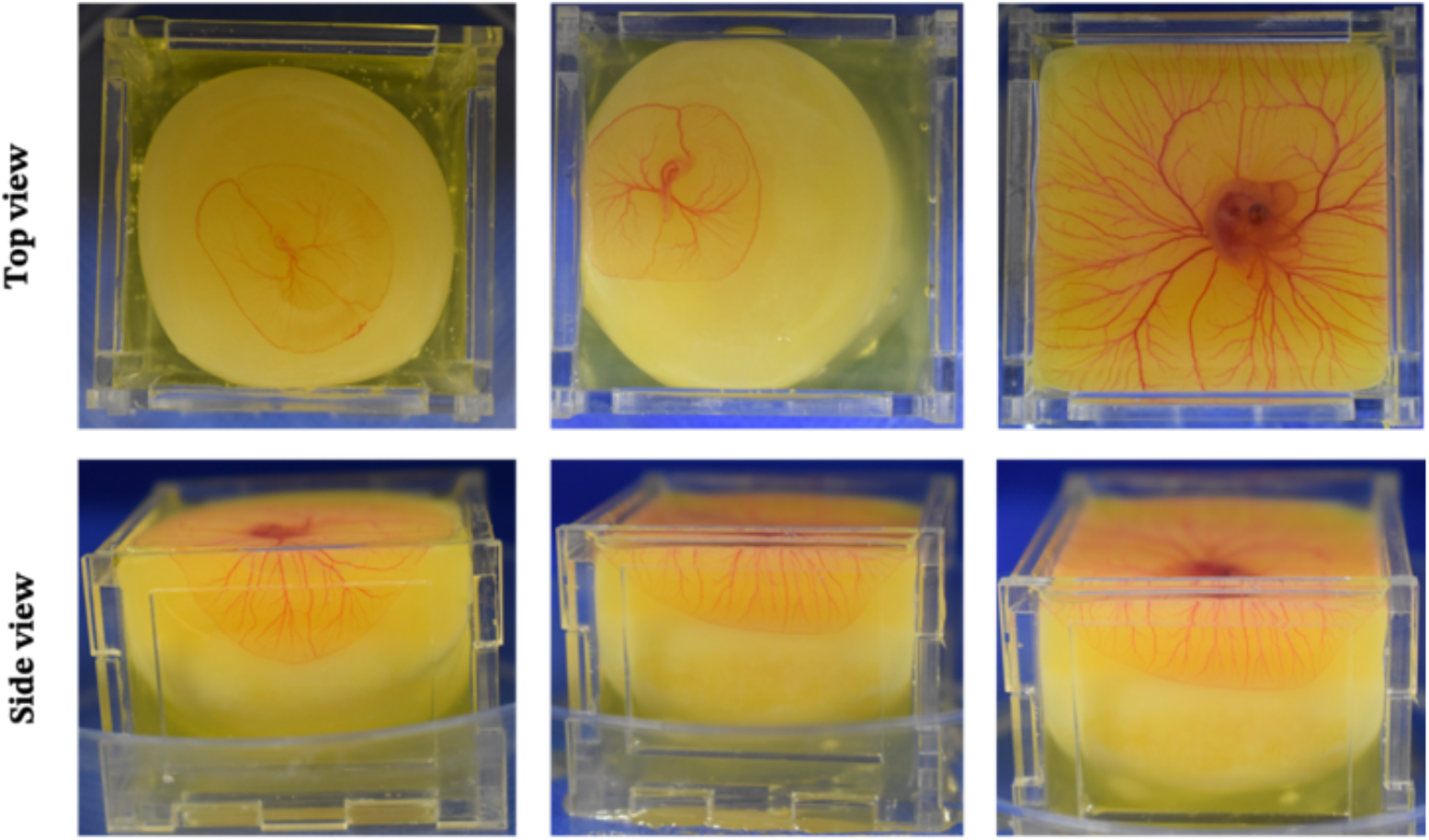
Fertilized egg showing CAM vasculature cultured within engineered eggshell.

**S3.**
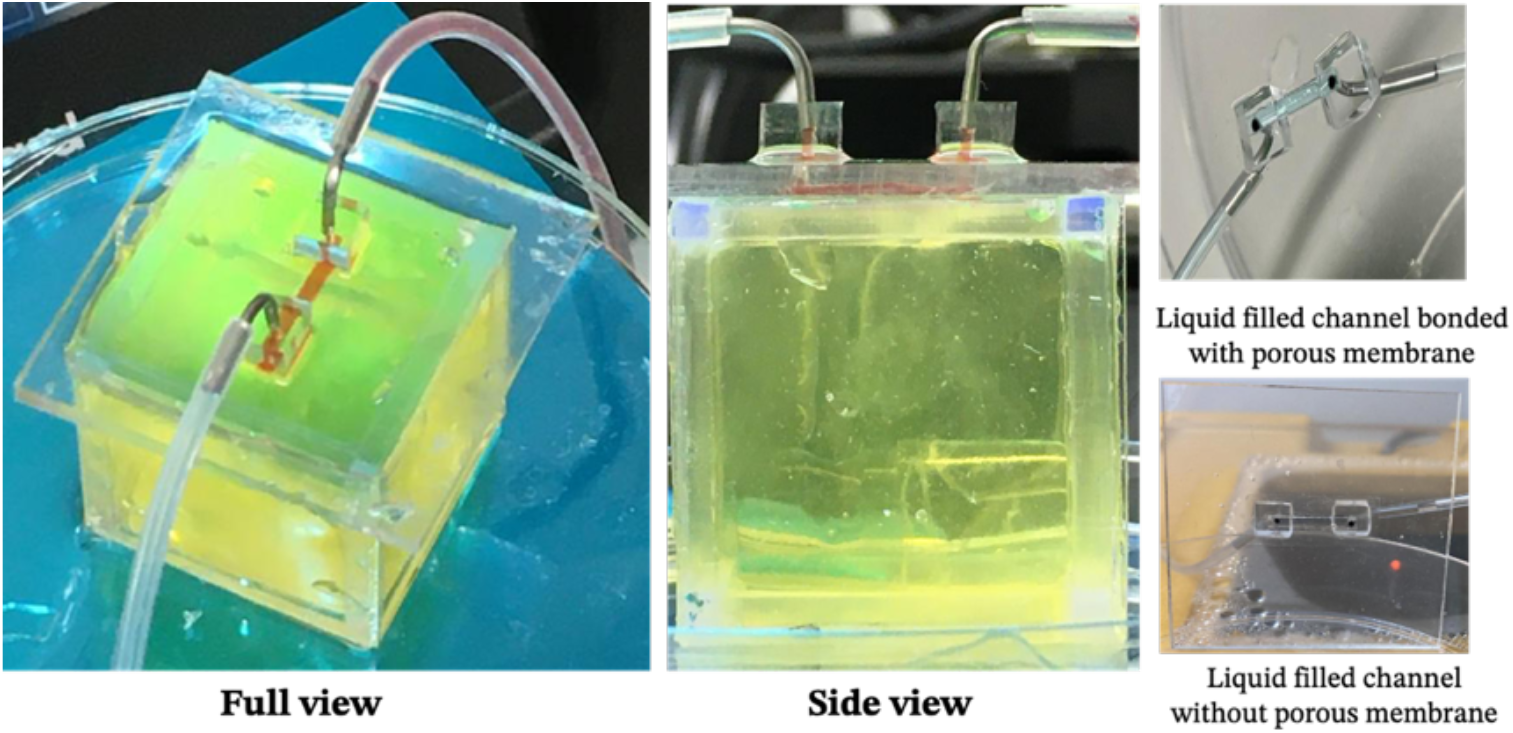
Proof-of-concept test: Impact of superfusion on porous membrane.

