## Supplementary Video Files S1-S5 for "Switching to external flows: perturbations of developing vasculature within chicken chorioallantoic membrane"

### Slide 1
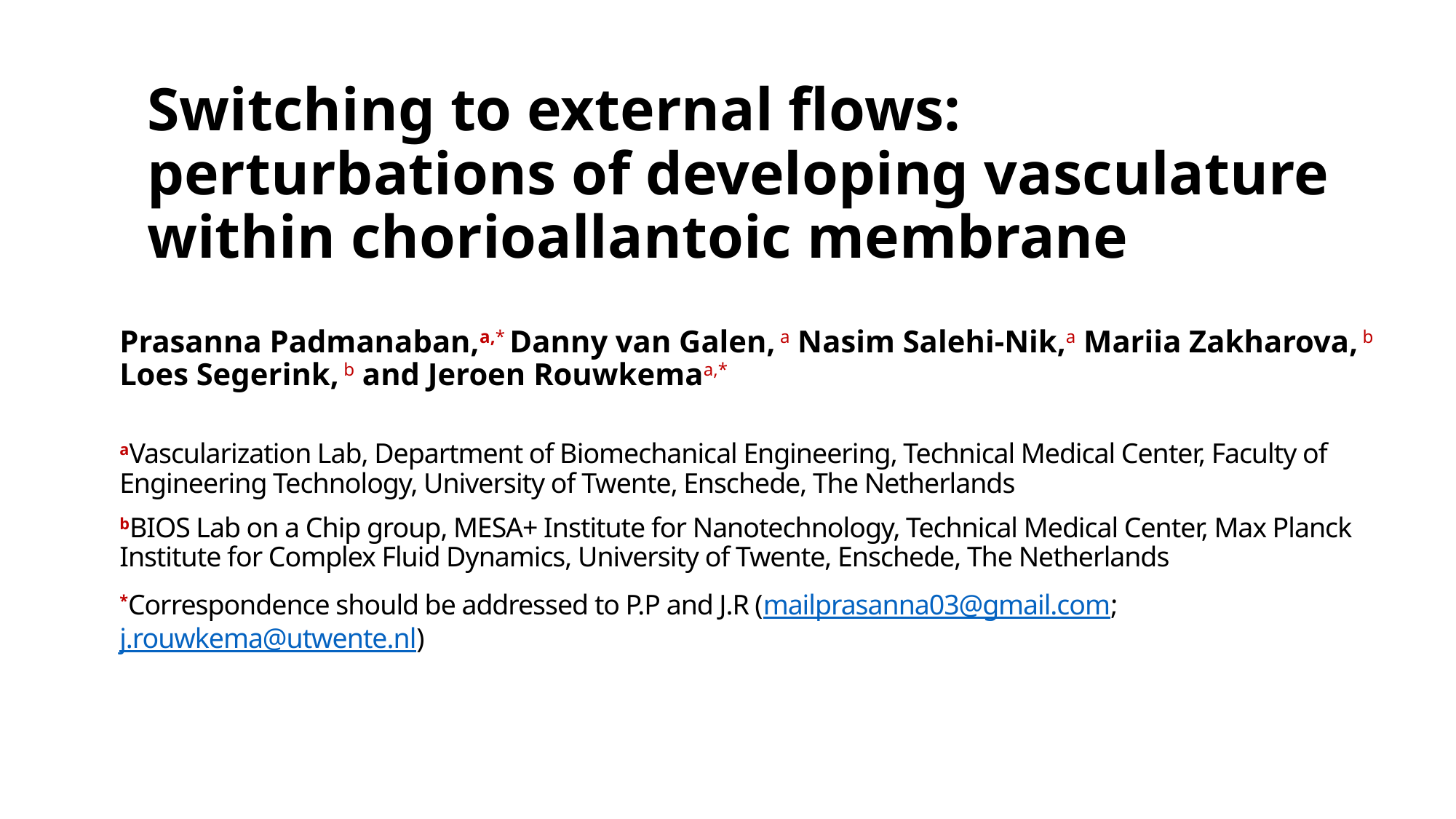

#### Switching to external flows: perturbations of developing vasculature within chorioallantoic membrane
Prasanna Padmanaban,^a,*^ Danny van Galen, ^a^ Nasim Salehi-Nik,^a^ Mariia Zakharova, ^b^ Loes Segerink, ^b^ and Jeroen Rouwkema^a,*^
^a^Vascularization Lab, Department of Biomechanical Engineering, Technical Medical Center, Faculty of Engineering Technology, University of Twente, Enschede, The Netherlands
^b^BIOS Lab on a Chip group, MESA+ Institute for Nanotechnology, Technical Medical Center, Max Planck Institute for Complex Fluid Dynamics, University of Twente, Enschede, The Netherlands

### Slide 2
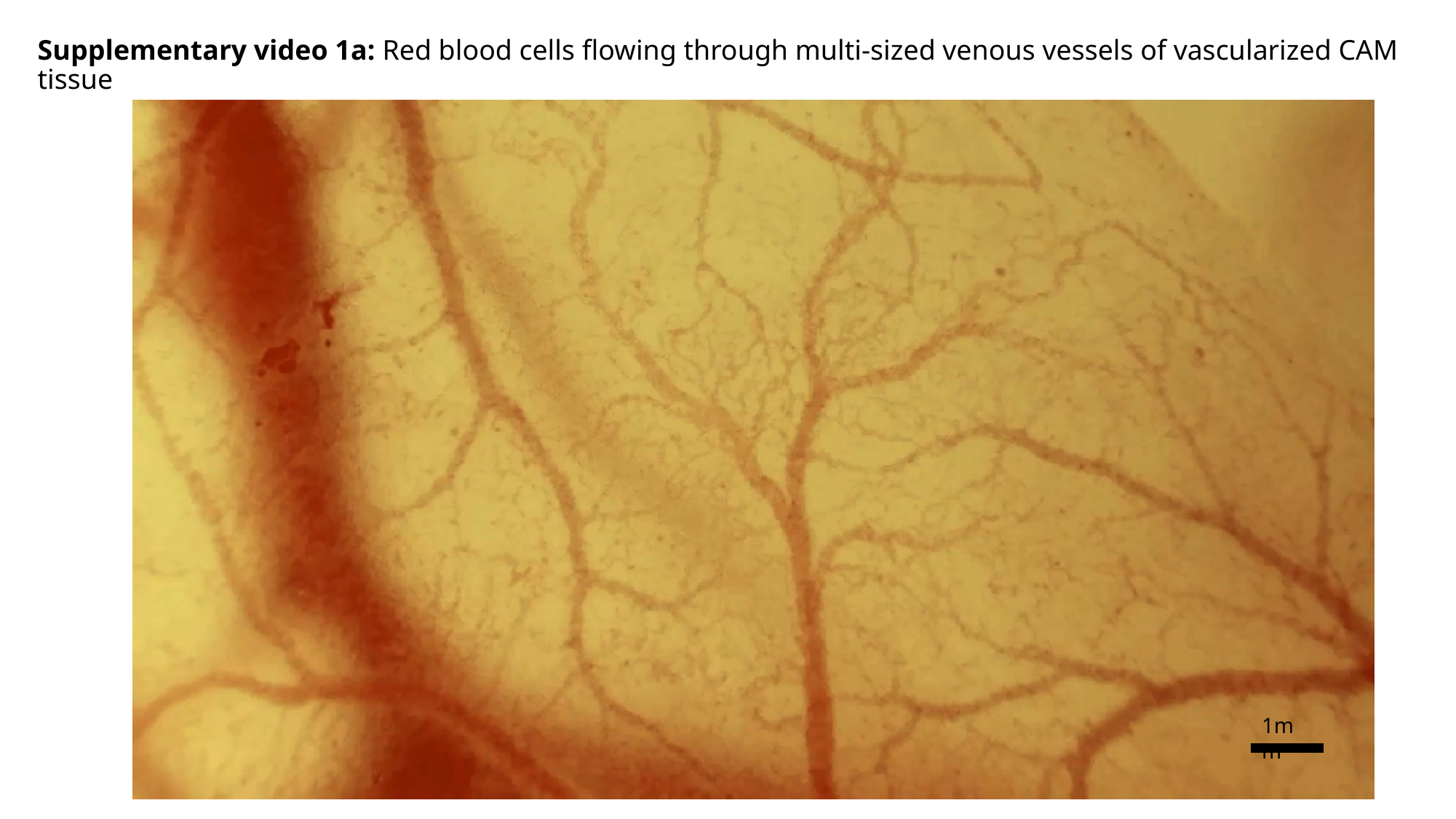

#### Supplementary video 1a: Red blood cells flowing through multi-sized venous vessels of vascularized CAM tissue
1mm

### Slide 3
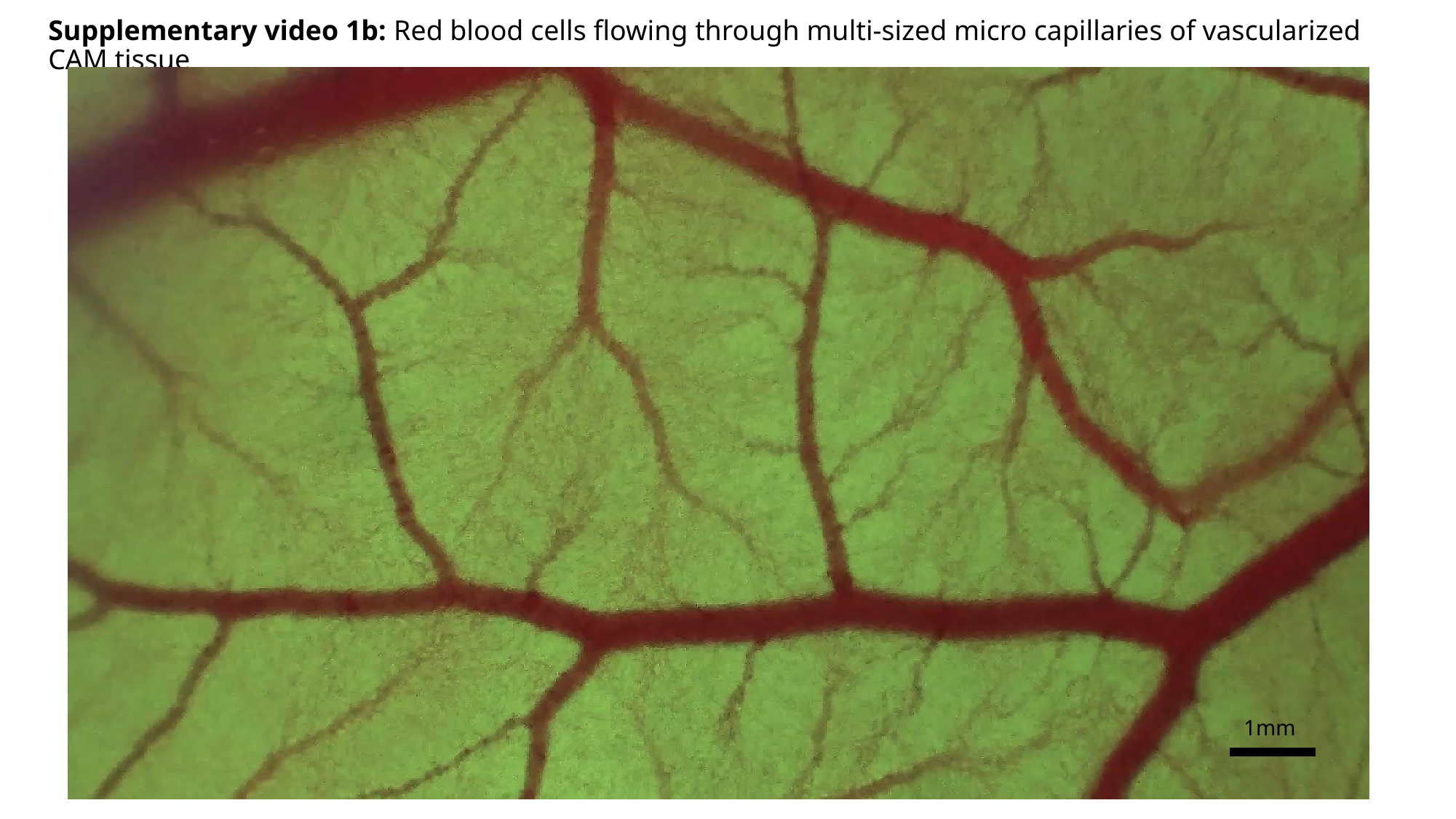

#### Supplementary video 1b: Red blood cells flowing through multi-sized micro capillaries of vascularized CAM tissue
1mm

### Slide 4
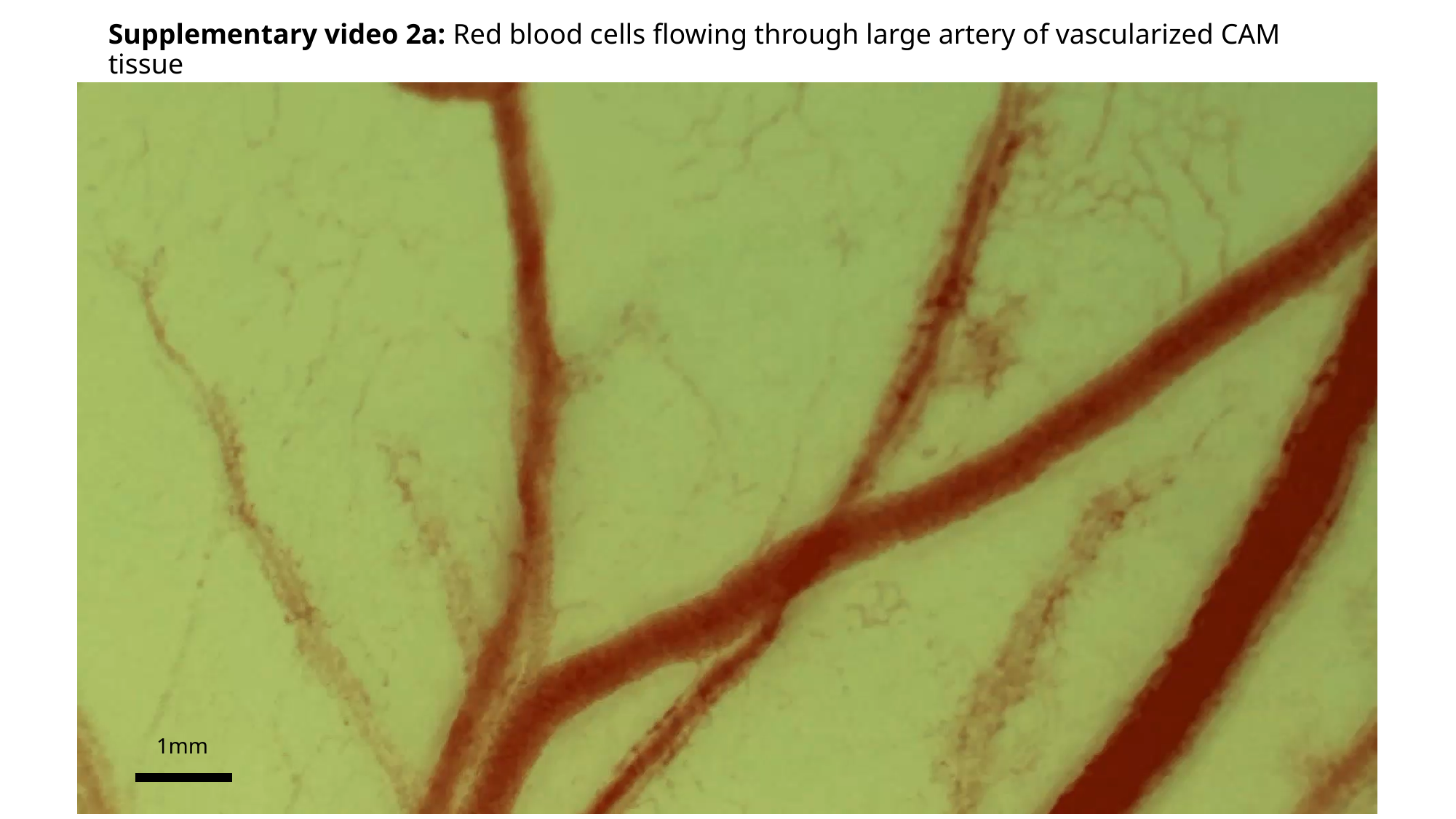

#### Supplementary video 2a: Red blood cells flowing through large artery of vascularized CAM tissue
1mm

### Slide 5
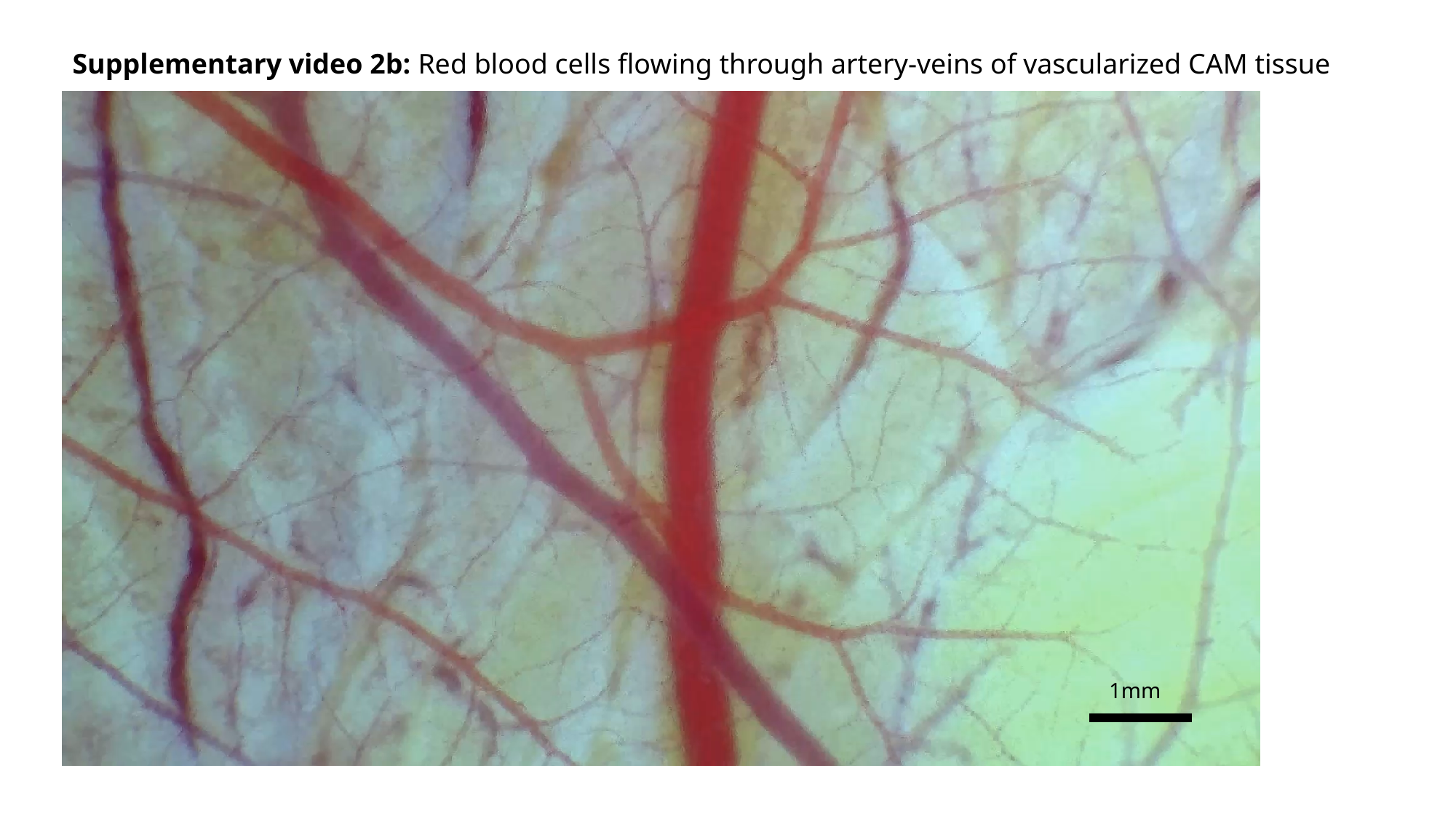

#### Supplementary video 2b: Red blood cells flowing through artery-veins of vascularized CAM tissue
1mm

### Slide 6
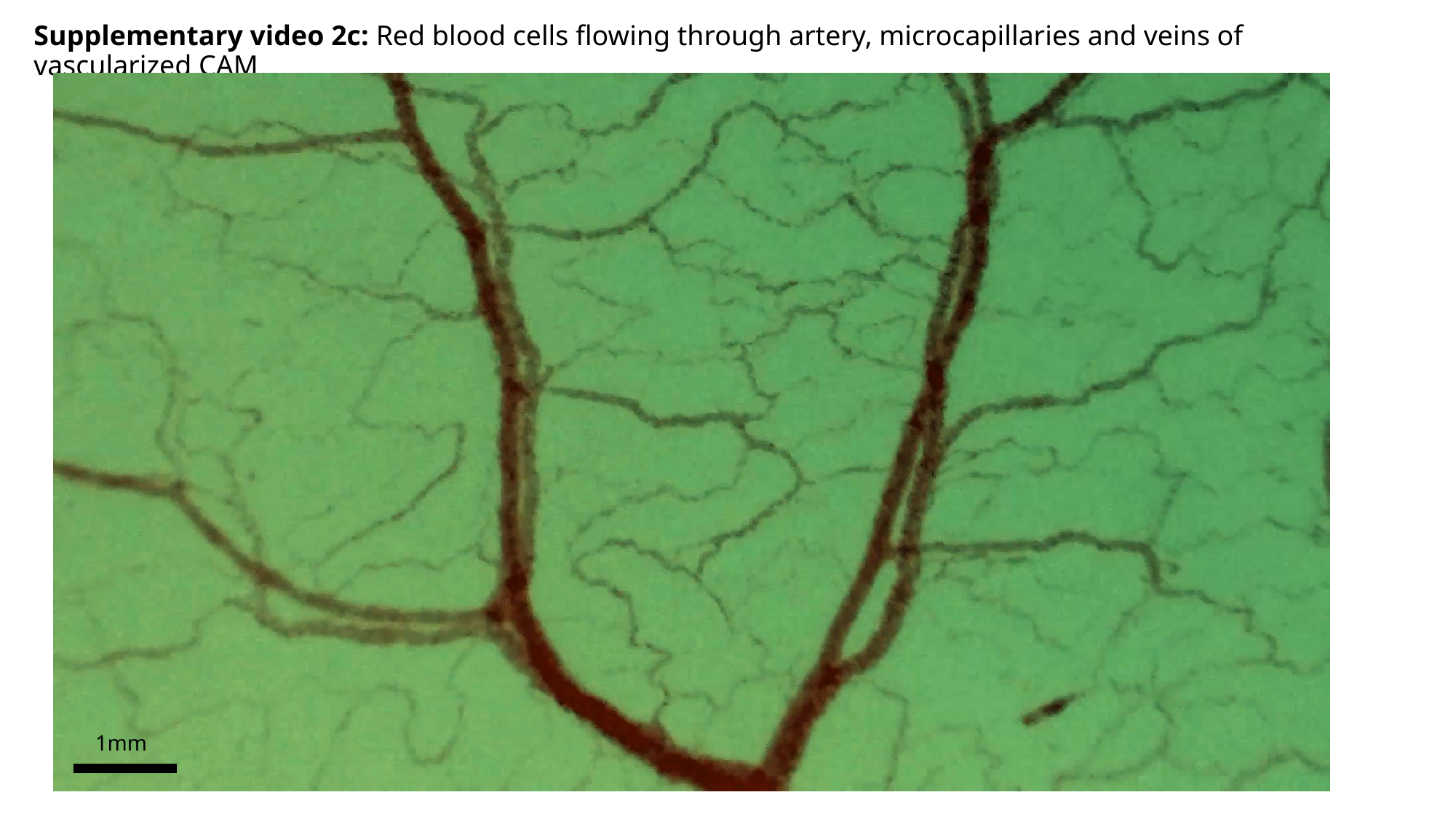

#### Supplementary video 2c: Red blood cells flowing through artery, microcapillaries and veins of vascularized CAM
1mm

### Slide 7
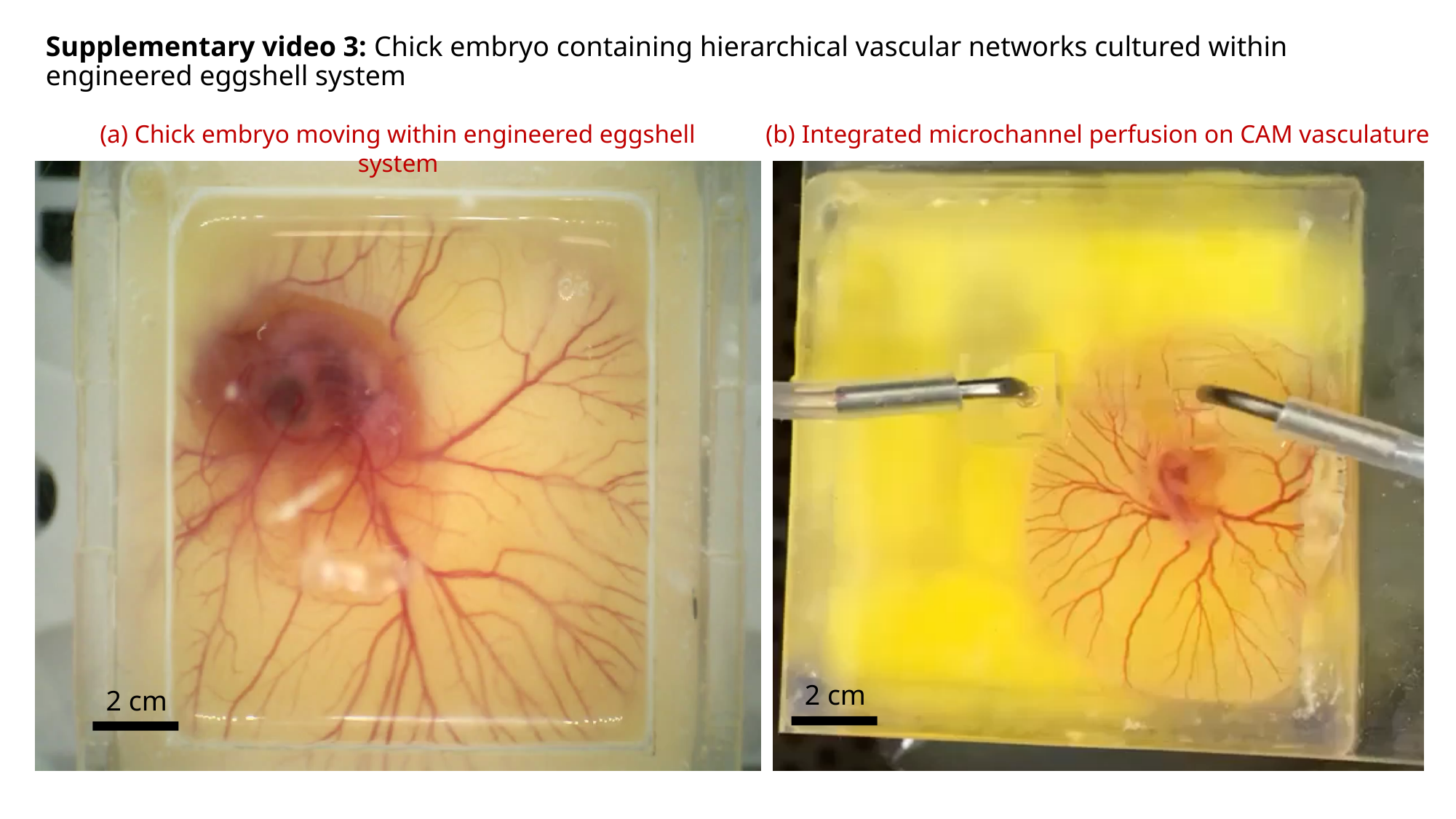

#### Supplementary video 3: Chick embryo containing hierarchical vascular networks cultured within engineered eggshell system
(a) Chick embryo moving within engineered eggshell system
2 cm
(b) Integrated microchannel perfusion on CAM vasculature
2 cm

### Slide 8
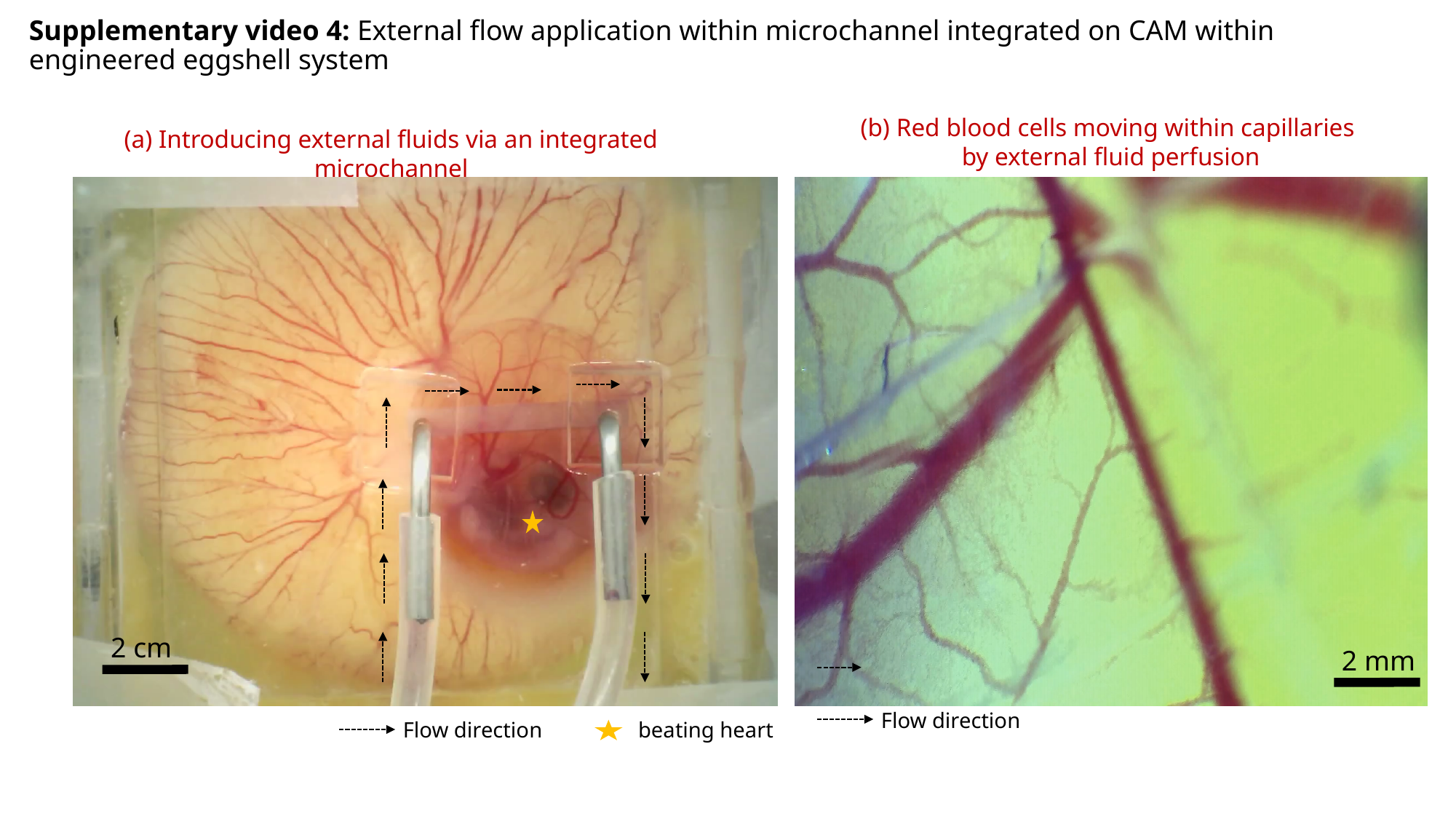

#### Supplementary video 4: External flow application within microchannel integrated on CAM within engineered eggshell system
(b) Red blood cells moving within capillaries
by external fluid perfusion
2 mm
Flow direction
(a) Introducing external fluids via an integrated microchannel
2 cm
Flow direction
beating heart

### Slide 9
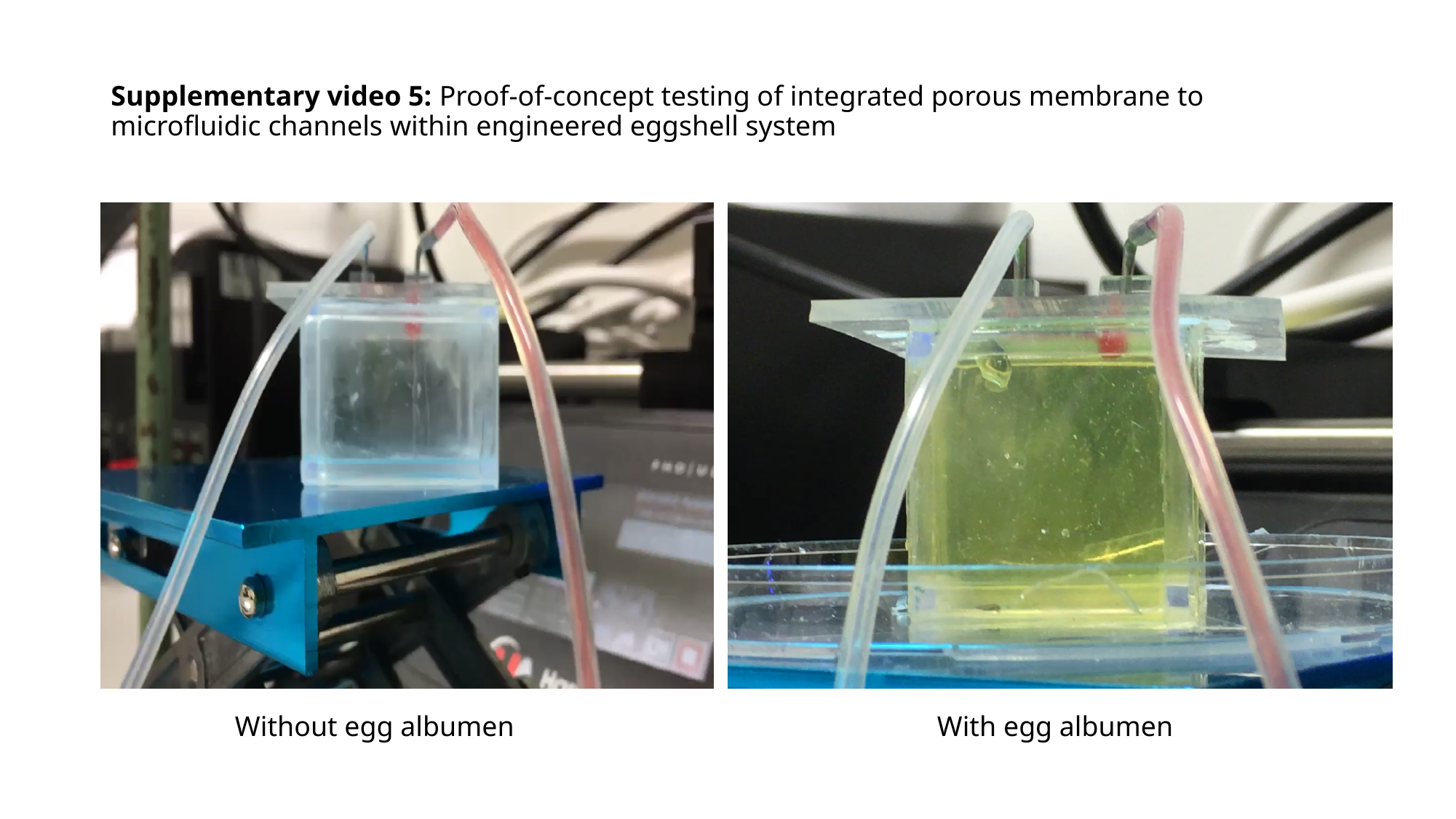

#### Supplementary video 5: Proof-of-concept testing of integrated porous membrane to microfluidic channels within engineered eggshell system
Without egg albumen
With egg albumen
